# Three-dimensional reconstruction of individual helical nano-filament structures from atomic force microscopy topographs

**DOI:** 10.1101/2020.02.28.970426

**Authors:** Liisa Lutter, Christopher Serpell, Mick Tuite, Louise Serpell, Wei-Feng Xue

## Abstract

Atomic force microscopy, AFM, is a powerful tool that can produce detailed topographical images of individual nano-structures with a high signal-to-noise ratio without the need for ensemble averaging. However, the application of AFM in structural biology has been hampered by the tip-sample convolution effect, which distorts images of nano-structures, particularly those that are of similar dimensions to the cantilever probe tips used in AFM. Here we show that the tip-sample convolution results in a feature-dependent and non-uniform distribution of image resolution on AFM topographs. We show how this effect can be utilised in structural studies of nano-sized upward convex objects such as spherical or filamentous molecular assemblies deposited on a flat surface, because it causes ‘magnification’ of such objects in AFM topographs. Subsequently, this enhancement effect is harnessed through contact-point based deconvolution of AFM topographs. Here, the application of this approach is demonstrated through the 3D reconstruction of the surface envelope of individual helical amyloid filaments without the need of cross-particle averaging using the contact-deconvoluted AFM topographs. Resolving the structural variations of individual macromolecular assemblies within inherently heterogeneous populations is paramount for mechanistic understanding of many biological phenomena such as amyloid toxicity and prion strains. The approach presented here will also facilitate the use of AFM for high-resolution structural studies and integrative structural biology analysis of single molecular assemblies.

## INTRODUCTION

Atomic force microscopy (AFM) is a scanning probe microscopy method that enables the collection of three-dimensional topographic image data, and has been widely applied to characterisation of biological and non-biological macromolecules. AFM encompasses a range of techniques for structural studies of biological molecules, including the study of inter- and intramolecular interactions [1], molecular dynamics [2], molecular remodelling under force [3] and imaging of molecules in air [4] or in liquid [5,6]. AFM operating in non-contact mode is capable of reaching atomic resolution on samples of small molecules [7], whereas AFM imaging of biomolecules routinely reaches nanometre resolutions in single high signal-to-noise images, and is able to characterise biological populations at a true single molecule level. This technique has been applied to a range of different bio-molecules, including membrane proteins [8], viral capsids [9] and filamentous biomolecules, such as amyloid fibrils [10], nucleic acids [11,12], and various filaments involved in the cytoskeleton [13].

Amyloid fibrils represent a class of filamentous supramolecular assemblies frequently studied using AFM imaging. They can be self-assembled from a wide range of different proteins and peptides. Fibrils formed from some amyloidogenic sequences, including amyloid-b and tau, have been identified to be involved in human neurodegenerative pathologies, such as Alzheimer’s disease, whereas other amyloid forming sequences are known to be non-toxic and even required for normal physiological processes [14]. Although the amino acid sequences of the many amyloid forming proteins are typically unrelated, the fibrils they form share a well-defined common core structural architecture, namely the cross-β arrangement made up of β-strands that stack perpendicular to the fibril axis stabilised by intermolecular hydrogen bonds between β-strands that runs parallel to the fibril axis [15,16]. Despite that all amyloid fibrils share these core structural features, variations in filament packing arrangements, including fibrils assembled from the same precursors, result in a multitude of different fibril structures called polymorphs, which may be related to the varying types of biological response they elicit. Furthermore, specific structural polymorphs assembled from the same amyloid forming protein could be patient- or disease-specific and may correlate with neurodegenerative disease aetiology [14]. Due to the unresolved nature of the amyloid assembly structure-function relationship, fibrillar amyloid specimen are widely studied using AFM imaging to resolve the mechanistic roles of amyloid assembly and polymorphism [17].

AFM imaging of a sample, e.g. amyloid fibrils deposited on a surface, with typical intermittent contact imaging mode such as the tapping mode is achieved by scanning a sample on a flat surface using a sharp tip attached to a cantilever. As the cantilever oscillates up and down, the tip and sample are moved relative to each other along the horizontal x and y axes in a raster pattern. When the tip interacts with the sample, vertical displacement of the cantilever relative to sample determines the surface height on the z-axis, measured at discrete pixel locations in the xy-plane. The lateral (x- and y-dimension) sampling is determined by the user in terms of the number of pixels collected per image and the pixel size, which tends to have edge length in the order of Ångströms to nanometres. The output of AFM scans is a 3-dimensional topography map, usually represented by a 2-dimensional coloured image where pixel values contain information on sample surface height, typically represented by the intensity of colour. Resolution in the vertical and lateral dimensions of AFM topographs are distinct. Topographs have a high vertical signal-to-noise ratio, limited by noise in the order of sub-Ångströms, due to electrical noise in the detector and thermal noise causing fluctuations in the cantilever [18,19], whereas lateral resolution is defined by the pixel size and the tip dimensions. Both vertical and lateral resolution can be affected by distortions or artefacts caused by drift, ambient noise or deformation of the sample or tip through their interactions. However, a significant limitation to the application of AFM to structural characterisation of biomolecules has been the tip-sample convolution effect, which causes lateral broadening, or dilation, of upwards convex structural features on AFM topographs. Convolution arises from the finite size and geometry of the AFM probe tip and is especially pronounced when sample features are of similar size to the tip radius, usually between 1-10 nm, as is the case for many biological macromolecules. The tip convolution effect has been studied in detail since the development of the AFM technique in the 1980s [20]. In order to minimise the convolution effect, both experimental and computational approaches have been developed. For example, various tip modifications have been used in order to minimise the effect of tip-sample convolution some of which have led to atomic resolution topographs [7]. However, this application is limited to small flat molecules and is not currently suitable for imaging biological macromolecules. The advantage of a computational approach to deconvolution is that an algorithm can be applied to a topograph of any specimen, after data collection. Many deconvolution methods have been reported to date. However common to these methods, they rely on the same concept based on the ‘erosion’ algorithm with various mathematical approaches [21–23]. Although erosion corrects for the dilation of the sample width, it leads to a significant loss of lateral structural information present in the AFM topology images because it does not recognise or recover the structural information that is present at subpixel locations, as we will show below.

In this report, we show that the AFM tip-sample convolution effect result in a feature-dependent and non-uniform distribution of image resolution on AFM topographs, and is in fact an advantage in terms of image resolution, as it causes the ‘magnification’ of the sample surface. Upon correction, we reveal that subpixel resolution information of the sample surface can be recovered using a contact-point based deconvolution algorithm, thus revealing the enhanced lateral resolution encoded in the AFM topographs while minimising the dilation image distortion. Furthermore, while AFM imaging provides only information on the top of 3-dimensional molecular surfaces of the sample structures, we show that helically symmetrical structures can be reconstructed as 3D surface envelopes from AFM topographs. This approach is demonstrated on imaging analysis of twisted amyloid fibrils formed from short peptide sequences which have been previously characterised using X-ray fibre diffraction [24]. The deconvolution and 3D modelling approach facilitates the structural analysis of individual twisted filamentous assemblies at an individual particle level. It is therefore capable of resolving the structural variations of individual macromolecular assemblies within inherently heterogeneous populations such as amyloid fibrils, which is key for mechanistic understanding of many biological phenomenon such as amyloid toxicity and prion strains.

## RESULTS

### Structural information is lost through erosion deconvolution of AFM topographs

The tip-sample convolution artefact arises from the finite shape and geometry of the cantilever tip. When the tip interacts with the sample, the surface height at the bottom of the tip is recorded, whereas the true surface of the sample may lie at a different location, probed at the tip-sample contact points (Figure 1a). This leads to convolution, or dilation in the case of upward convex surface features on AFM topographs. Image deconvolution by the erosion method corrects for the dilation of the sample by translating a model of the tip to the coordinates at which surface heights were recorded on the image and finding the tip’s deepest penetration of the convoluted surface to reconstruct a deconvoluted image while conserving pixel coordinates and sampling in the xy-plane (Figure 1b and 1c) [23,25]. Thus, erosion resolves apparent clashes between the tip and convoluted sample surface image by lowering sample surface heights according to the known surface values of the tip translates. AFM tip geometry can be typically approximated as conical with varying apical radii, and side angles that can be modelled from the values provided by the manufacturer. Erosion deconvolution is efficient in minimising the dilating effect of convolution. However, it causes a significant loss of structural information (demonstrated on a twisted amyloid fibril in Figure 1c), because it replaces the existing vertical surface height values with the surface of deepest tip penetration without finer resampling of surface heights at tip-sample contact points, which contain structural information in the images. These contact points lie off the pixel grid at subpixel locations and contain structural information on the true sample surface. As shown on the twisted amyloid fibril example (Figure 1), deconvolution by erosion causes loss of information on the fibril helical periodicity and molecular surface features.

**Figure 1.**
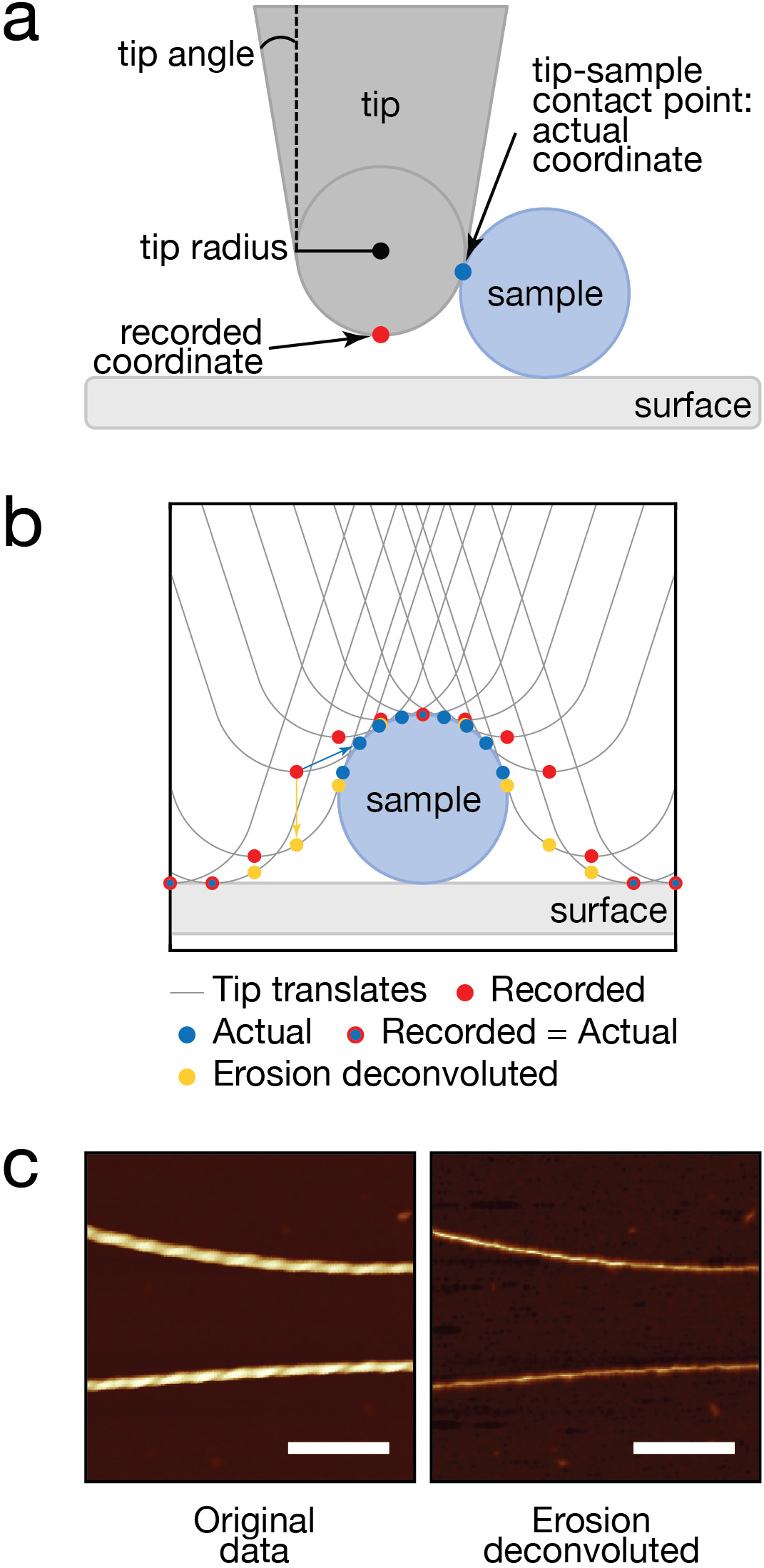
AFM image deconvolution by the erosion algorithm corrects for dilation but causes significant loss of structural information of the sample surface. **(a)** Schematic cross-sectional illustration of AFM tip-sample interactions that lead to lateral convolution and dilation of sample features. This imaging artefact is especially pronounced for sample objects on the same size range as the tip radius. **(b)** Schematic cross-sectional diagram of tip-sample deconvolution by erosion. The sample cross-section is illustrated as a blue circle. Red dots show original coordinates of the data, orange dots show erosion deconvoluted height coordinates at each pixel location and blue dots show the coordinates at tip-sample contact points for each pixel location. **(c)** Example of deconvolution by the erosion algorithm demonstrated on an example image of a twisted amyloid fibril. Left is an AFM topology image showing twisted amyloid fibrils. Right image shows the erosion-deconvoluted image of the same data, demonstrating substantial loss of structural information due to the erosion algorithm. A symmetric conical tip with a side angle of 18° (estimated from the tip geometry information provided by the manufacturer) and tip radius of 11.2 nm (estimated using the image features as described in the Results) was used as a model of the tip. The scale bars represent 200 nm.

### Tip-sample convolution results in enhanced lateral sampling

Although the tip-sample convolution effect causes upward convex surface features to appear dilated on topographs, this effect can in fact be use to advantage as it causes the ‘magnification’ of structural features present at tip-sample contact points located at subpixel coordinates, and results in higher lateral resolution topographic information to be captured than what is recorded at pixel grid coordinates. This information can be recovered by a deconvolution method in which the tip, with a known geometry, radius and angle, is modelled on the recorded surface scan at translations that correspond to original lateral sampling (x and y coordinates) and recorded surface heights (z coordinates), similarly to the erosion deconvolution method. However, instead of finding the surface of deepest penetration, geometric modelling of tip and simulation of tip translates is used to find tip-sample contact points. These tip-sample contact points contain information on the true surface of the sample, which is otherwise recorded in magnified, or dilated, form. Thus, the total number of data points remains the same but the pixel coordinates in both xy-plane and surface heights on the z-axis, are shifted, revealing the enhanced local sampling of the upward convex surface features encoded in the image data. This contact point deconvolution approach can be applied to any AFM topograph and is demonstrated on a sphere, a cylinder, and a randomly generated rough surface (Figure 2). Contact points are found by iterative rounds of tip-sample simulations, in which the sample surface is initially assumed to be circular, until convergence of the deconvoluted surface coordinates. The corrected images can then be interpolated onto a finely spaced even grid for visualisation of the deconvoluted image. In figure 2, lateral sampling of the tops of the example objects and peaks of the random rough surface can be seen to increase in density and shift significantly from the original position of xy-gridlines (Figure 2, 3rd column), especially for features with size similar to the radius of the tip.

**Figure 2.**
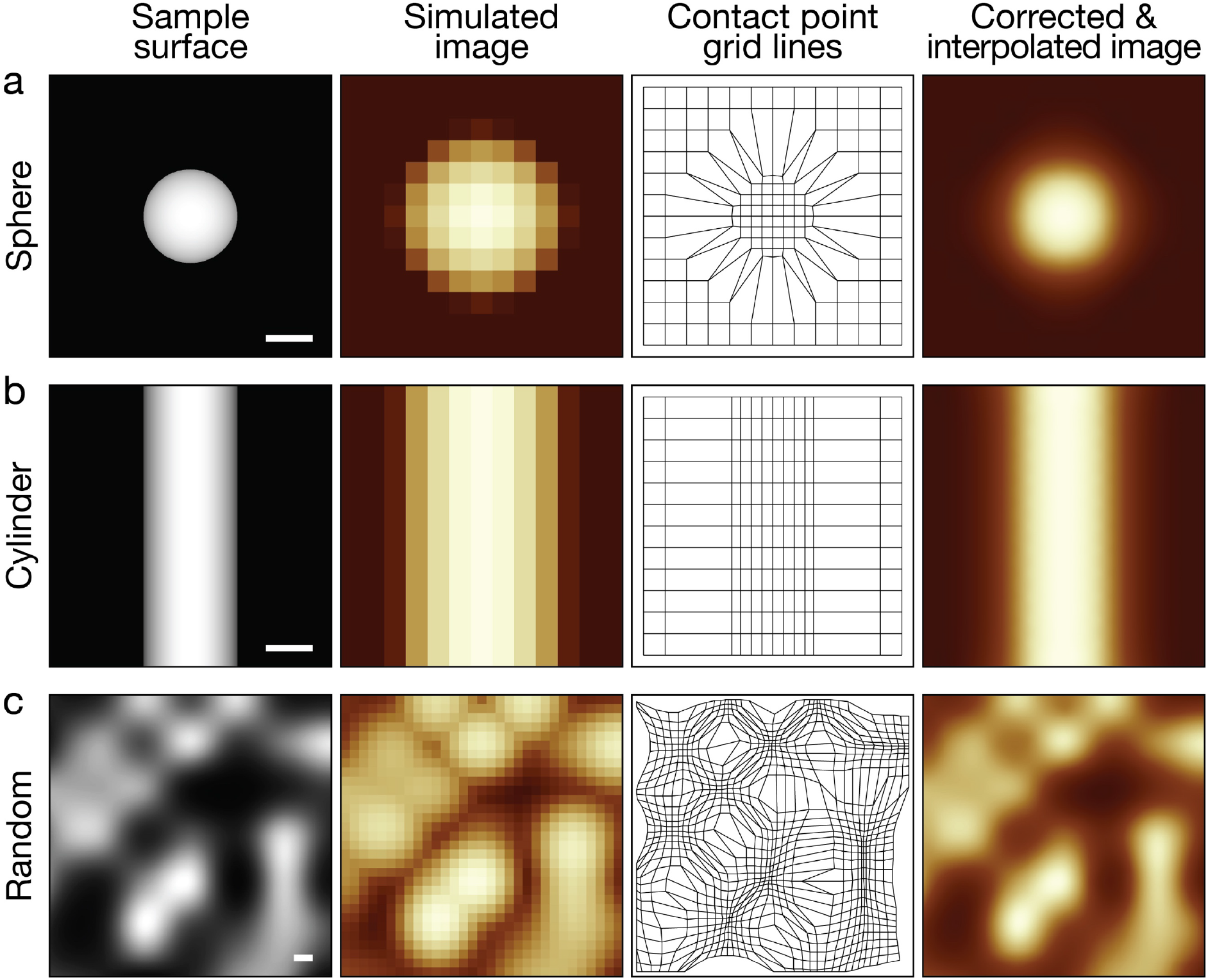
Contact-point deconvolution can be applied to any surface and results in increased local lateral sampling for upward convex sample features. Simulated AFM image (2nd column from left), shifted pixel grid lines after contact point deconvolution (3rd column from left) and a corrected image with contact points interpolated to a finer pixel grid (right most column) are shown for a sphere **(a)**, cylinder **(b)**, and a random surface **(c)** shown in the left-most column. The sphere and the cylinder both have a radius of 2 nm. A symmetric conical tip with a radius of 2 nm and tip side angle of 18° was used for all simulations. The scale bars represent 2 nm.

The effect of locating the contact points off the pixel grid on the images of upward concave features is, however, less favourable as the depth of a trough can only be accessed if the sides of the tip do not come into contact with the sample. This can be seen in the Fig 2 bottom row example as the decrease in contact point density in these areas. In the case of biological filaments such as DNA, cytoskeletal and amyloid filaments, major upward concave surface features include grooves, which can vary ~1-50 nm in width. They may be resolvable by AFM, depending on the geometric parameters of the tip, as well as depth of the features.

The deconvolution and lateral sampling enhancement of a convoluted surface depends on both the geometry of the sample, geometry of the tip, and the lateral pixel sampling frequency. The effect of these various factors on the sampling enhancement is here assessed with scan-line simulations using a circular cross-section as the sample and a symmetric tip model constructed from typical tip parameters (Figure 3). The sampling enhancement factor is measured as a ratio of deconvoluted image signal density to original image signal density, where signal density is found as the number of moved pixel coordinates that result from tip-sample contact per surface area. The sampling enhancement factor describes the increase in lateral sampling frequency as a result of tip-sample contact point deconvolution. The effect of sample diameter as well as tip radius on the enhancement factor is illustrated in Figure 3. As seen, the sampling enhancement factor of a sub-nm circular cross-section is more than 3 with a 2 nm tip-radius, representing more than 3 times increase in lateral sampling compared to the convoluted image (Figure 3 left column). The sampling enhancement then decreases as the circle diameter increases and plateaus at ~1.4 times enhancement compared to a convoluted surface. The sampling enhancement fluctuates for non-continuous sampling realistic to actual imaging experiments and the stepwise large increases represent increases at which the tip and circle first come into contact at a sampled pixel as the radius of the circle increases, allowing a new contact point to be found. Enhancement then slightly decreases with the increase in circle radius as no new contact points are added and existing contact points become spaced further apart, therefore decreasing information density.

**Figure 3.**
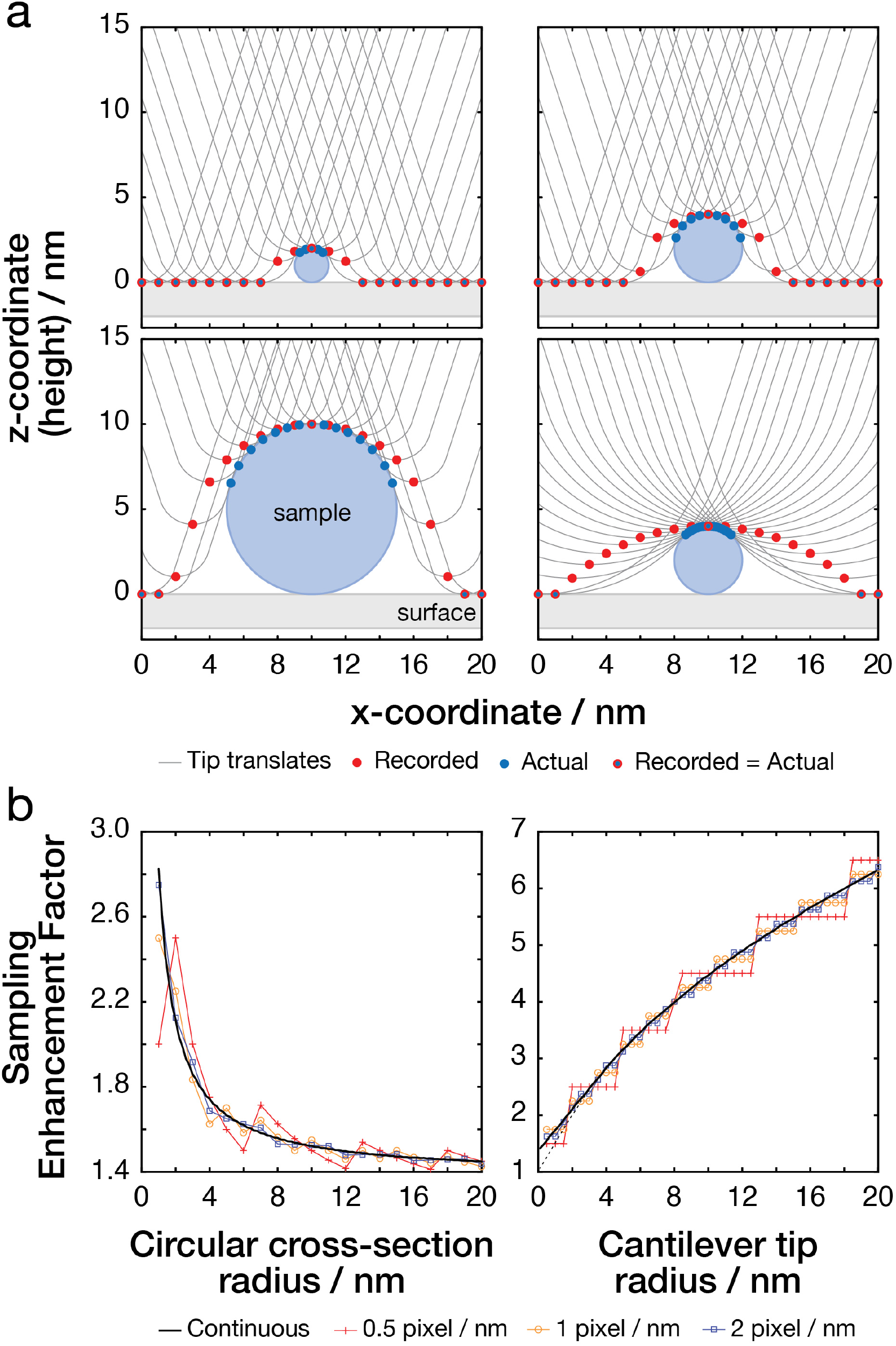
Image sampling enhancement effect after image correction by contact-point deconvolution shown with varying sample and sampling parameters. Schematic diagrams are shown in (a) and results from simulations in (b). Red circles on schematics represent original convoluted coordinates at each sampled pixel and blue circles represent tip-sample contact points at each sampled pixel. The sampling enhancement factor is defined as the ratio of pixel density of these points within the circle envelope. Left column shows the enhancement effect seen with a circle with varying radius. A symmetric conical tip model with a radius of 2 nm and tip side-angle of 18° was used. Right column shows the enhancement effect observed with a circle with a constant radius of 2 nm with varying tip radius and a tip side-angle of 18 °. The dashed line represents the same simulation but with a tip with side-angle of 0°.

For a symmetric conical tip, the most important tip parameters for determining tip-sample interactions are the apical radius of the tip and the angle at which the sides of the tip widen. Although AFM tips with smaller tip radii and tip angles produce higher resolution images by minimising the tip-sample convolution effect, with the contact-point deconvolution method, larger tip radii and tip angles result in higher sampling enhancement as the tip interacts with the sample at more xy-coordinates, allowing proportionally more contact points to be recovered (Figure 3b). Thus, with finite scanning precision, increasing resolution of images becomes an optimisation problem that involves finding optimum tip geometry with any given sampling frequency and specimen geometry. Importantly, simulations of tip-sample interactions with an estimate of sample geometry or an already corrected sample surface in which tip model and pixel size parameters are varied can guide the selection of an optimal tip geometry and sampling frequency to maximise resulting image resolution for a specific sample.

### Contact-point deconvolution of an AFM amyloid fibril topograph

The deconvolution algorithm and resulting lateral sampling enhancement are demonstrated on an AFM topograph of an amyloid fibril formed from a short amyloid forming peptide with the amino acid sequence HYFNIF (Materials and Methods and [17]. The fibril example is first traced from the AFM image across the fibril central line and subsequently straightened and interpolated to an evenly spaced pixel grid [26], while maintaining pixel size identical to that in raw data. In order to apply the contact point deconvolution algorithm as shown on the examples above, the convoluted surface, as well as a model of the tip used to scan the specimen are needed. For experimental image data, the variation in the tip radius from their nominal value, which results from the tip manufacturing process, should be considered [27]. It is also important to consider that the tip can become blunter with scanning and its tip radius can widen over time. Here, the tip radius can be estimated for each individual fibril on an image from the extent of convolution seen in data. An estimate of the tip radius is found by assuming the twisted amyloid fibrils have ideal corkscrew symmetry, and the average cross-section of the fibril perpendicular to its axis of rotation is, therefore, circular with a radius defined by half of the maximal z-height value. Least-square regression analysis is then performed to fit a simulated convoluted scan-line generated from modelling interactions of the tip with the circular cross-section model to the average convoluted cross-section observed in data, while letting the tip radius vary as a parameter. It is assumed that the overall tip geometry and side angles do not change. In the example shown in Fig 4, while the nominal tip radius provided by the manufacturer was 2 nm, using the approach described here the tip radius estimate was 11.2 nm. The deconvolution algorithm is then applied to the 3D topograph to find corrected grid lines and surface heights (Figure 4c,d). The deconvoluted contact points follow the twisting pattern of the fibril and increase lateral sampling of the fibril surface as predicted. The ungridded data points are then interpolated back onto a finer evenly spaced grid for visualisation of the de-convoluted contact-points (Figure 4c).

**Figure 4.**
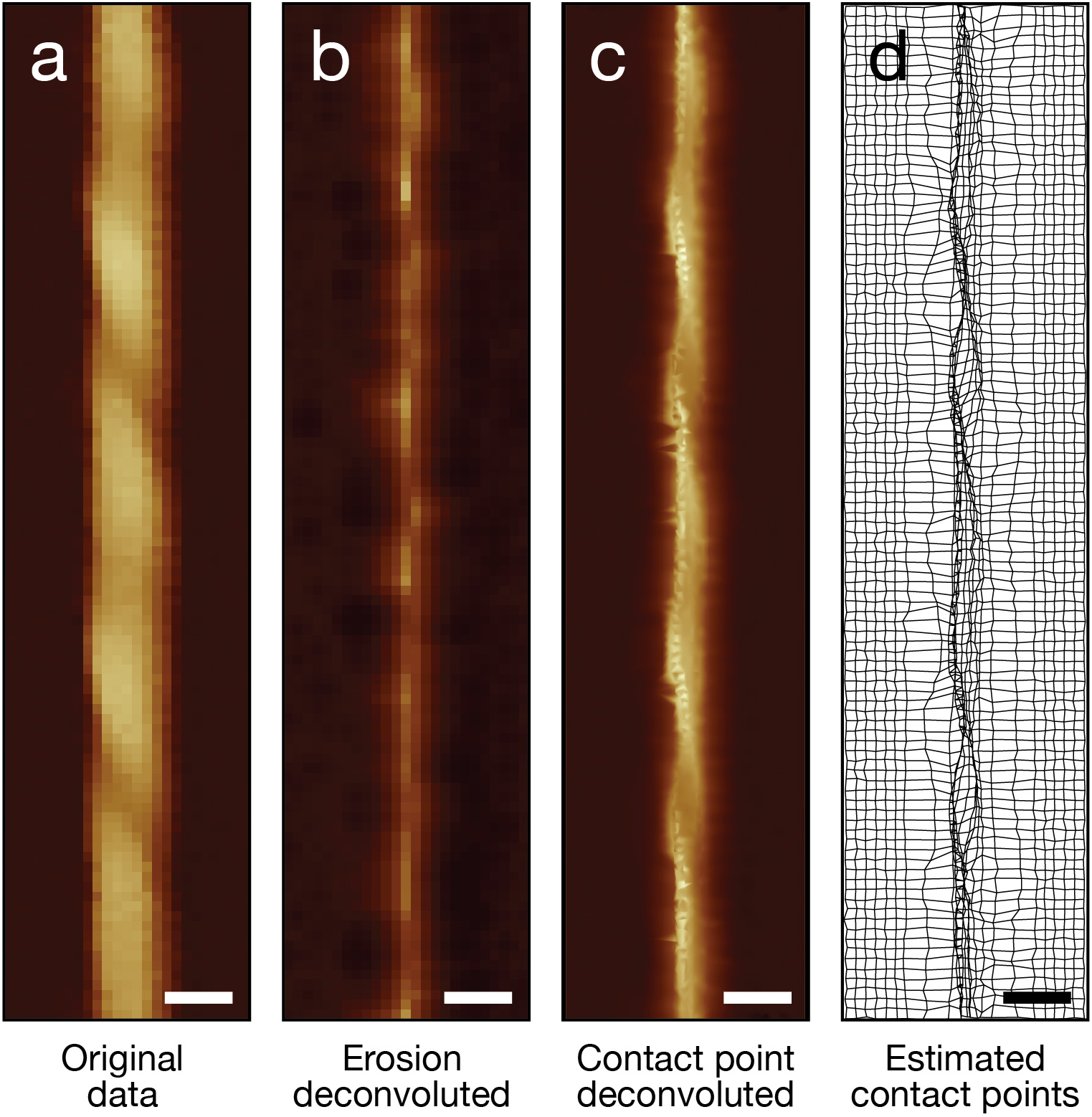
Typical example of contact-point deconvolution of an AFM topograph of a helically twisted amyloid fibril. (**a)** Digitally straightened AFM topography image of an amyloid fibril. **(b)** Image deconvoluted using the erosion algorithm. **(c)** Contact-point deconvoluted and interpolated image. (d) Corrected lateral (x/y) grid lines after tip-sample contact-point deconvolution. A symmetric conical tip model with a radius of 11.2 nm and side angle of 18° was used. A short section of the total fibril is shown for detail and the corresponding part of the same fibril is shown on each panel. The scale bars represent 20 nm.

### Assessment of AFM topograph lateral resolution

Because AFM topographs are extraordinarily high signal-to-noise, the lateral resolution of a single uncorrected AFM topographs is typically determined by the spatial frequency of sampling in the xy-plane. According to the Nyquist sampling theorem, the minimal sampling rate that contains the information to reconstruct a signal is twice the maximum frequency component, thus the Nyquist resolution limit for images is twice the pixel size. For example, the image used in the example in Figure 4 has a pixel size of 2.93 nm and the resolution is, therefore, limited to 5.86 nm. Deconvolution using our approach shifts pixel coordinates to subpixel positions off the pixel grid. Therefore, recovering structural information of the sample by shifting the sample surface coordinates present in magnified form results in recovering of the true higher lateral sampling frequencies present in the image data. Figure 5 shows the lateral resolution of deconvoluted AFM images assessed using a feature-based lateral resolution assessment method [28] and compared with the original convoluted image of the same fibril shown in Figure 4. The feature-based method applies a low-pass filter to an image in spatial frequency domain from higher towards lower spatial frequencies and measures the correlation between the filtered and unfiltered images in real space. Increases in correlation, seen as peaks in a log-log plot of the first derivative of cross-correlation vs. spatial frequencies, indicate presence of structural information at the specific spatial frequency. Figure 5b shows the correlation curves with peaks indicating the presence of structural information. Comparison of the normalised correlation between the convoluted and deconvoluted amyloid fibril images shows a shift of the correlation curve towards the right, indicating a shift of information content towards higher spatial resolutions due to correcting of sampling frequency during deconvolution. However, the contact point deconvolution has a much larger effect on the x-axis (across the width of the filament) than on the y-axis (along the length of the filament), which leads to broadening of the peaks on the 1^st^ derivative of correlation graph. Analysis of lateral resolution suggests that the highest resolution at which structural information is found on the convoluted example fibril image is ~60 Å (consistent with the Nyquist frequency of the original image data) and on the deconvoluted image the highest resolution is ~30 Å, indicating that deconvolution results in approximately doubling of the lateral resolution, for this amyloid fibril, due to the recovery of the enhanced sampling in the image data.

**Figure 5.**
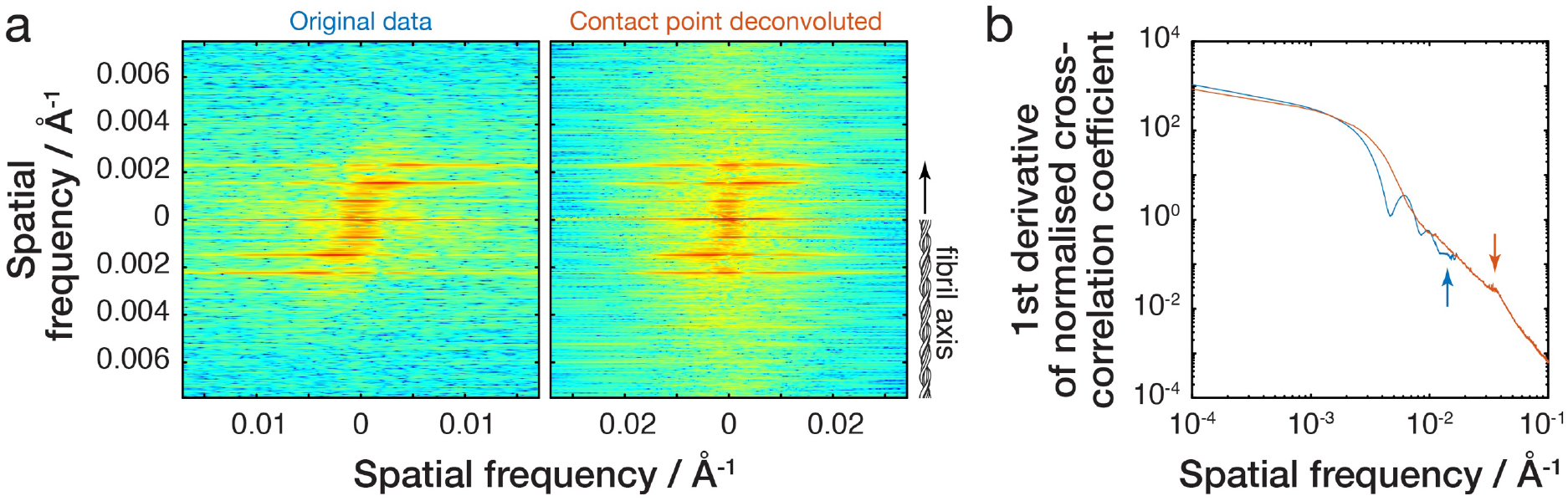
Resolution of original recorded and corrected AFM topographs are evaluated by the feature-based lateral resolution assessment method [28]. The same fibril image as Figure 4 is used as example. **(a)** 2D Fourier spectra of the original convoluted (left) and contact-point deconvoluted (right) images. The x-scale is expanded to accommodate peaks in the spectra for contact-point deconvoluted image as they are shifted to higher spatial frequencies. **(b)** Log-log plot of the first derivative of the normalised cross-correlation curve. Arrows indicate right-most peaks indicating the highest spatial frequency information present at 64Å resolution for the original convoluted fibril image and 28Å for the contact-point deconvoluted image.

### 3D modelling of helical amyloid fibrils from corrected AFM topographs

While AFM provides 3-dimensional topographic data, only the top surface of the sample is accessible to the tip, preventing the visualisation of the full sample surface envelopes in 3D. However, for helically symmetric structures such as amyloid fibrils, we describe below an algorithm that can be used to reconstruct 3D surface envelopes by taking advantage of the screw-axis symmetry of the filaments. This approach is demonstrated on AFM topographs of deconvoluted amyloid fibrils. The workflow of AFM image processing for 3D surface envelope reconstructions is summarised below (Table 1).

**Table 1:**
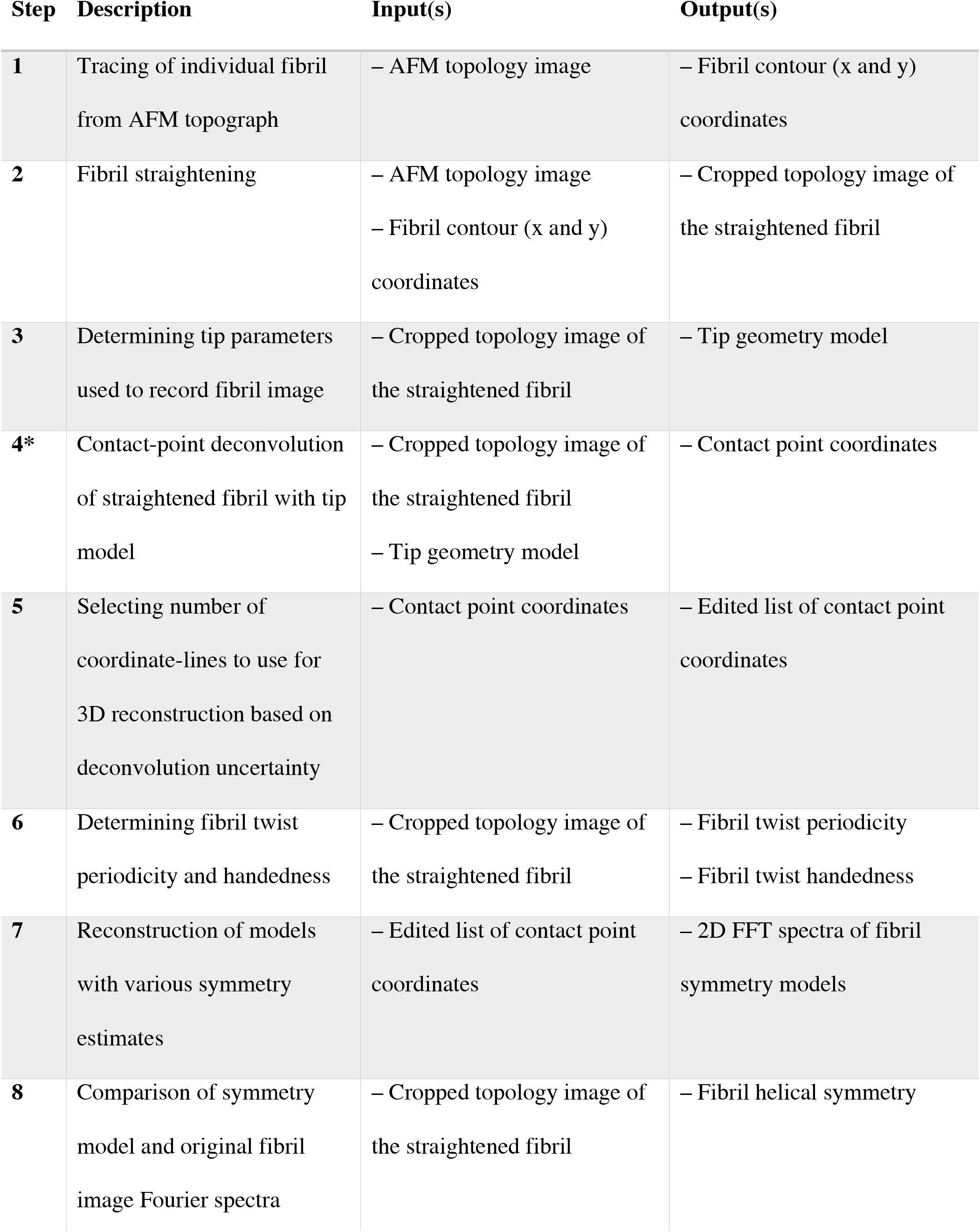
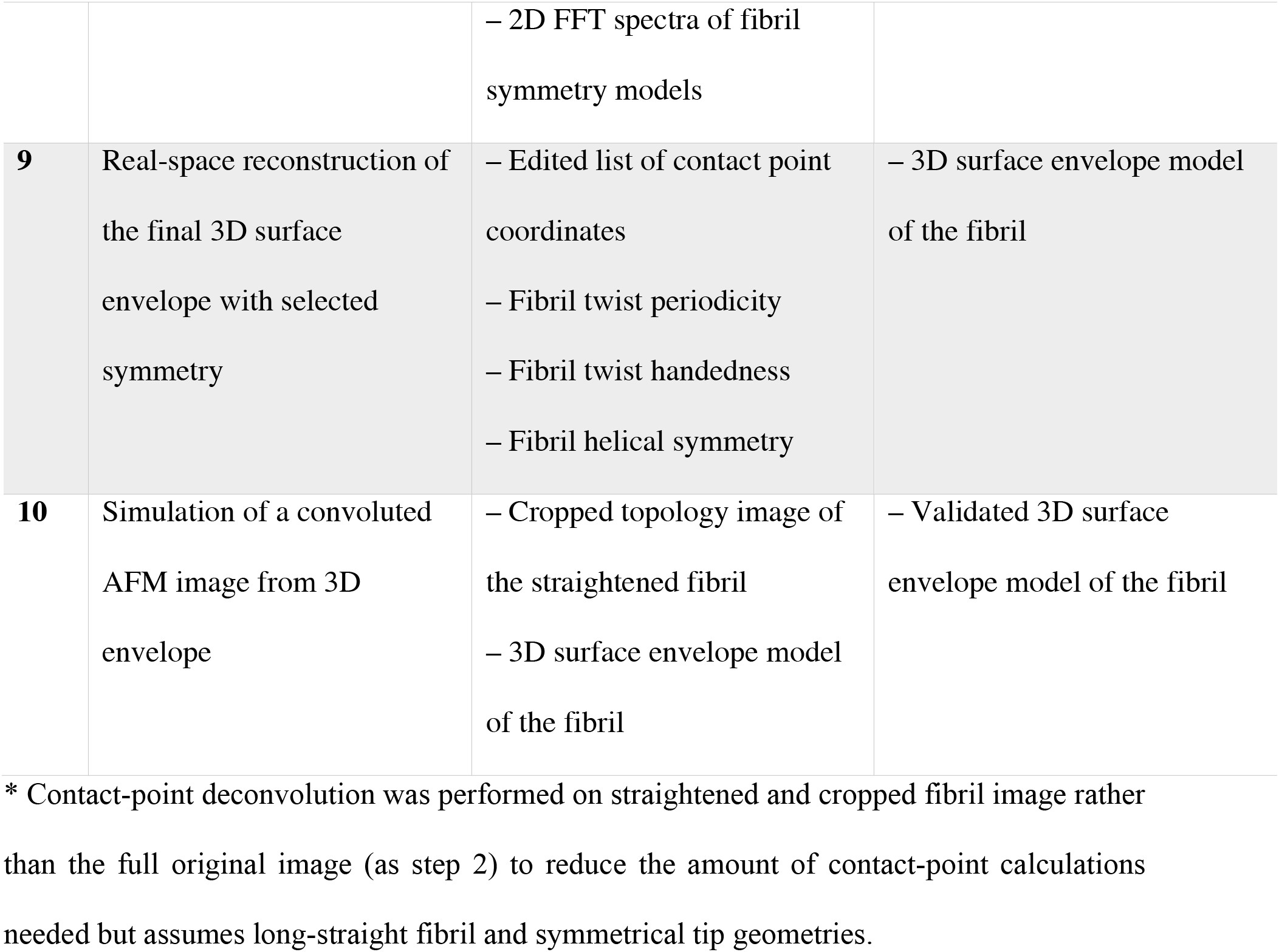
Flow chart of AFM topograph processing and 3D helical fibril surface envelope reconstruction.

Following contact-point deconvolution of traced and digitally straightened fibril image, the surface envelopes are reconstructed using a moving window approach in which rotation and translation is applied to each corrected contact point coordinates within the window, thus reconstructing fibril cross-sections as the window is slid along the length of the fibril. The width of the moving window is one complete rotation of the fibril cross-section along the fibril axis (i.e. one nominal helical pitch), determined by its periodicity and screw axis symmetry. The rotation of each data point within the window depends on the distance of the point from the central cross-section of the window. Rotation angle values are negative in one direction from the centre of the window and positive in the other direction, with the specific direction depending on the twist-handedness of the fibril. The rotation angles are kept constant along the length of the filament. After deconvolution of the amyloid fibril, the number of lines along the length of the filament that are to be used for the 3D reconstruction are determined. Data points further away from the centre of the filament can be more frequently affected by artefacts during scanning and the uncertainty of the deconvolution algorithm tends to be higher for surface heights that lie further away from the central line. However, data points further away from the centre of the filament also contain additional structural information and information on the fibril twist and width which is harder to determine from the lines closer to the centre alone. Therefore, the number of pixel-lines used is determined by visual inspection of the deconvolution result. The deconvoluted surface coordinates that correspond to the selected pixel-lines are then used for the 3D modelling approach. The deconvoluted contact points off the pixel-grid are interpolated onto an evenly spaced grid along the screw axis, creating ‘slices’ perpendicular to the fibril axis, while the data remains ungridded along the x-axis. In order to determine the length of the reconstruction window for the deconvoluted amyloid fibril image, the periodicity of the fibril is first determined by applying a 1D Fast Fourier Transform (FFT) to the surface height profile of the fibril central line (Figure 6a-b). This determines the spatial frequencies of repeating patterns in the signal. The spatial frequency with the highest amplitude represents the periodicity of the fibril. For an asymmetric fibril cross-sections, one period represents a complete turn (one helical pitch) whereas for a fibril with *n*-fold symmetry periodicity represents *1/n* of a complete turn. Fibril symmetry is then estimated by constructing 3D models with systematically varied symmetries and comparing these to the original straightened fibril image to find the best match (Figure 6c). Symmetry determination is a critical step for reconstructing the surface of a helical specimen. Different screw axis symmetries result in differences in the twisting pattern observed on the fibril top surface. In order to make the comparison of a 2D convoluted image and 3D deconvoluted reconstructions, the tip model is used to simulate convoluted AFM images from the symmetry models. The straightened experimental fibril image and various symmetry simulations are zero-padded to square images and compared as 2D Fourier spectra to estimate fibril screw-axis symmetry that best describes the original image data. The 2D Fourier spectra contain information on repeating patterns of fibril top surface and are analogous to a diffraction pattern. Fourier-based analysis of helical biomolecule structures dates to first diffractions patterns that describe the structure of DNA and has been since used to reconstruct helical biomolecules from electron cryo-microscopy images by indexing the Bessel functions of the diffraction pattern layer lines [29]. However, in many cases, resolving individual layer lines and indexing them is not possible, especially for amyloid fibrils with long helical repeat distances and small twist angles of individual subunits. Differences in rotational symmetry of a fibril cross-section causes the Bessel orders to change significantly while the layer lines on the spectrum remain identical [30]. Here, using this property and assuming that the screw-axis of the filament lies straight along the centre of the fibril, the differences in layer line Bessel orders observed for different symmetry models, which contain information on the handedness and screw axis symmetry of the fibril and represent the twisting pattern on top of the fibril, were used to estimate the fibril screw-axis symmetry. For example, if the line between most intense off-centre spots in the 2D FFT (+ symbols in Figure 6c) has a positive slope then the fibril is left-handed, and if the slope is negative the fibril is right-handed. The steepness of the slope contains information on the screw-axis symmetry of the fibril and the closest match of a symmetry model to the original fibril image is used to estimate the symmetry for an individual fibril. For the amyloid fibril example shown in this demonstration (Figure 6), the 2-fold symmetry model gives the best fit to data (Figure 6c), suggesting that the fibril cross-section has a pseudo 2-fold screw axis symmetry. These 2D Fourier spectra from AFM images are less ambiguous compared to analogous 2D Fourier patterns of TEM images as only the top of the fibril surface contributes to the signal, facilitating the direct measurement of handedness and symmetry.

**Figure 6.**
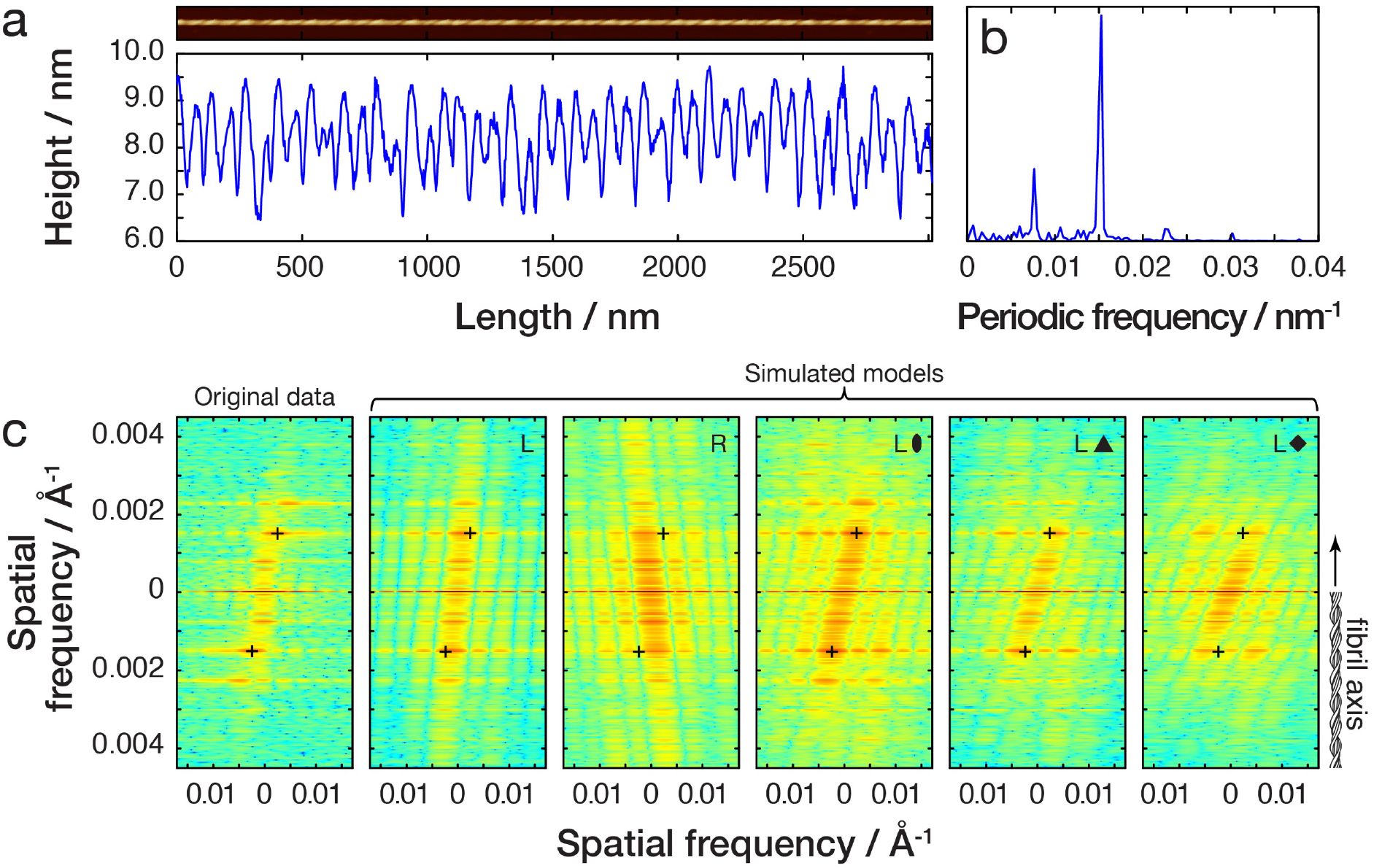
Estimation of the fibril twist periodicity and handedness. The same fibril image as Figure 4 is used as example. **(a)** Fibril central line height profile. **(b)** One-dimensional FFT power spectrum of the central line is used to determine the periodicity of the fibril. **(c)** The screw axis symmetry and the twist handedness of the fibril is estimated by reconstructing 3D models of the fibril with varying symmetries, simulating convoluted AFM images from the models and comparing the 2D FFT spectra of the symmetry model images to that of the original straightened fibril image. The crosses are drawn on the most intense peaks on the original fibril 2D FFT image and their position is then applied onto each symmetry model 2D FFT spectra to guide finding the closest match. In this example, a left-hand twisted fibril with a pseudo two-fold screw-axis symmetry shows the best match to the data.

Having determined the periodicity, handedness and screw-axis symmetry of the fibril, the 3D surface envelope can be reconstructed using the moving window approach described above (Figure 7). A cross-section of the fibril is obtained at each y-coordinate and a cubic spline is fitted to the cross-sections for smoothing. The number of spline pieces is determined by manual testing, taking into account the periodicity and symmetry of the fibril, which affects the number of times each cross-section is sampled. For validation of the image deconvolution and reconstructed 3D models, the final 3D surface envelope and tip model are used to simulate a convoluted AFM image that can then be compared with the uncorrected straightened fibril image (Figure 7b). Although there are small differences, the simulated image twist, periodicity and local surface features generally correspond well to the original fibril image, validating the reconstruction approach here as a useful method for reconstruction of individual fibrils without cross-particle averaging.

**Figure 7.**
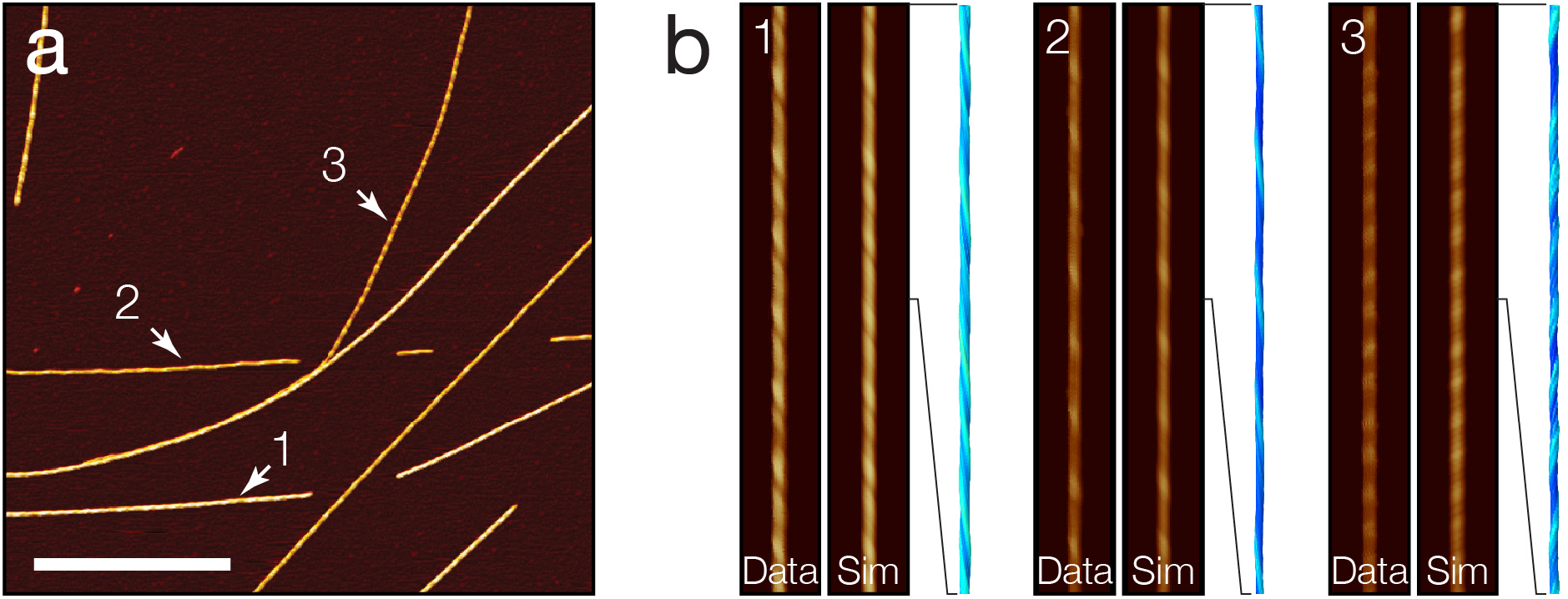
3D surface envelope reconstruction of amyloid fibrils from AFM topography. **(a)** AFM topology image of amyloid fibrils assembled from short peptide with the sequence HYFNIF [17,24]. The scale bars represent 1 μm. (b) 3D surface envelope models for each fibril indicated in (a) are shown together with a comparison of the original straightened uncorrected fibril images and the simulated AFM images from the final 3D models. Segments of 500 nm are shown for each fibril model and 1 μm for the images for detail.

## DISCUSSION

Tip-sample convolution leads to lateral dilation of upward convex surface features on AFM topographs, which has been seen as a significant limitation to the usefulness of AFM images for structural biology applications. Here we show that the convolution effect results in the magnification of the sample top surface and non-uniform distribution of structural information on the image. The structural information, present in an un-gridded form at sub-pixel tip-sample contact points, can be recovered at its true lateral resolutions without loss of information using a contact-point based deconvolution algorithm, which is demonstrated here on an amyloid fibril formed from a short peptide sequence. This approach improves on the previous erosion-based deconvolution algorithms and facilitates the use of AFM imaging of nanostructures in structural studies. Furthermore, we show how the 3-dimensional coordinates encoded in the topographic data on AFM images and the helical symmetry of the twisted fibrils can be used to reconstruct 3D surface envelopes, allowing analysis of structural parameters, such as the fibril cross-sectional area, which are not otherwise present directly on AFM images of these samples. Thus, the reconstruction will facilitate the integration of 3D AFM envelope models with data from other structural biology tools for integrated global structural analysis. The moving window approach of 3D surface reconstruction conserves the high signal-to-noise feature of single-molecule imaging capability of the AFM, thus also allowing the intra-fibrillar and local structural variation to be preserved in the 3D model. Deconvolution and modelling facilitate the application of AFM to characterise and quantify amyloid fibril polymorphism at a true individual single molecule level.

Although there are many techniques available for studying the structures of biological macromolecules, each with their own advantages and disadvantages, AFM occupies a unique position among these tools due to its high signal-to-noise ratio capable of resolving morphological features of individual molecules. Ensemble averaging of structural information from many molecules is a key concept on which techniques like X-ray crystallography and cryo-EM rely and has led to the elucidation of numerous atomic and near-atomic resolution structural models of biological macromolecules. For example, recent advances in cryo-EM equipment and data processing have allowed the reconstruction of protein structures in various conformations from a set of micrographs and can even contain information on molecular dynamics [31]. However, due to the low signal-to-noise ratio of individual particles on cryo-EM images, only ensemble-average structures can be reconstructed. Although the overall resolution of AFM images of macromolecular assemblies does not yet reach the order of Ångströms and only molecular surface features can be probed, it is able to resolve morphological features at the level of individual molecules. This allows AFM to tackle biological problems in a unique way, especially in cases where polymorphism and structural variations of molecules is important for their biological effects. The lateral resolution, as defined by pixel size and tip radii, tends to be around 1-5 nm, although vertical resolution can reach the order of sub-Ångströms. Using the contact-point deconvolution approach demonstrated here on AFM images of biological nano-filaments, recovery of the true lateral sampling resolution at tip-sample contact points was estimated to typically result in doubling of lateral resolution for helical nano-filaments. Experimental improvements, such as using a tip with parameters that maximise sampling enhancement and better scanners could lead to a more pronounced improvement in lateral resolution. Further improvements to the deconvolution and envelope reconstruction algorithm approach will also likewise help to improve the overall resolution achievable with AFM.

The structural basis of why amyloid fibrils can have functional roles in a wide range of organisms, associated with pathological symptoms of neurodegeneration, or simply exist as inert aggregates is not clear [14]. Individual filament 3D reconstruction using the approach presented here on AFM topographs could help elucidate the link between morphological features of the fibrils to specific biological responses of amyloid populations. Reconstructed 3D surface envelopes of amyloid fibrils could also be used in an integrative way with other structural biology techniques. For example, in recent years numerous cryo-EM reconstructions of amyloid filaments from *ex vivo* patient tissue have been resolved. These are ensemble averages from thousands or more of individual fibrils from the total fibril population, which in some cases may appear homogenous, but in other cases may be made up of a varying amount of fibril polymorphs. The number of some of these polymorphs within the population may also be too low for reconstruction, although these species might have some specific biological effect. 3D envelope reconstruction from AFM images could be used as a complementary technique as it allows the structural analysis and surface modelling of each individual fibril in the population, allowing rare members of the ensemble population as well as the population characteristics itself to be analysed. Classifying and quantifying the fibril structures could be used to determine the landscape of possible fibril structures within a specific population. Furthermore, this technique allows the intrafibrillar variation to be analysed, which may be indicative of fibril dynamics and stability. The 3D surface envelope could also be used in an integrative way with other techniques, contributing to the global information for modelling the atomic structure of a biomolecule.

The physics of AFM imaging is unique in producing its high signal-to-noise data at nano-scale and enables true single molecule approaches. Thus, the contact-point deconvolution and 3D envelope reconstruction approach presented here will facilitate the use of AFM for single-molecule structural studies. Imaging of any nano-structures with the AFM can benefit from the contact-point deconvolution approach to correct images while recovering structural information present at higher lateral resolutions. This includes high-speed AFM [32], which compromises spatial resolution for higher temporal resolution and allows imaging of biomolecule dynamics in the timescale down to milliseconds, and which could benefit from the computational deconvolution of image frames to recover the higher spatial resolution sampling of the sample surface. Furthermore, the 3D envelope reconstruction algorithm could also be applied to diverse biological samples with symmetries, including DNA, membrane proteins that form tubular arrays e.g. nicotinic acetylcholine receptor pore [33] and the mitochondrial outer membrane protein TspO [34], as well as helical or spherical virus capsids and cytoskeletal filaments. AFM imaging of various small molecules and polymers assembled into helical supramolecular arrangements which are widely used in chemistry, materials science and nanotechnology could, likewise, benefit from 3D surface reconstructions as AFM is widely used to characterise a wide range of such structures [35], for example polymer wrapped functionalised carbon nanotubes [36,37], Further developments could also lead to samples exhibiting different symmetries such as icosahedral viral capsids to be reconstructed. In conclusion, the approach reported here will facilitate the use of AFM for structural studies of individual molecules in complex populations, and for integrative structural biology analysis of single molecular assemblies and their assembly mechanisms.

## MATERIALS AND METHODS

### Peptide amyloid fibril synthesis

The amyloidogenic peptide HYFNIF was synthesised by N-terminal acetylation and a C-terminal amidation. Multistage solid phase synthesis using Fmoc protection chemistry was used to generate a lyophilised powder with > 95% purity measured by HPLC (JPT peptide technologies, or Biomolecular analysis facility, University of Kent). The lyophilised powder was suspended in 100 μl of filter sterilized milli-Q water to a final concentration of 10 mg/ml. The solution was incubated at room temperature for 1 week prior to imaging.

### AFM sample preparation and image acquisition

The peptide samples were diluted to 0.05 mg/ml in a solution of HCl (pH2, using filter sterilised milli-Q water). 20 μl of sample was deposited onto freshly cleaved mica (Agar scientific, F7013) and incubated for 10 minutes. Following incubation, the sample was washed with 1 ml of filter sterilised milli-Q water and then dried using a stream of nitrogen gas. Fibrils were imaged using a Multimode AFM with a Nanoscope V (Bruker) controller operating under peak force tapping mode with ScanAsyst probes (silicon nitride triangular tip with tip height = 2.5-2.8 μm, nominal tip radius = 2 nm, nominal spring constant 0.4 N/m, Bruker). Images were collected with a scan size of 6 x 6 μm with 2048 x 2048 pixel resolution. A scan rate of 0.305 Hz was used with a noise threshold of 0.5 nm and the Z limit was reduced to 1.5 μm. The peak force set point was set automatically, typically to ~675 pN during image acquisition. Nanoscope analysis software (Version 1.5, Bruker) were used to process the image data by flattening the height topology data to remove tilt and scanner bow.

### Image data analysis

Fibrils were traced and digitally straightened [26,38,39] using an in-house application and the height profile for each fibril was extracted from the centre contour line of the straightened fibrils. The periodicity of the fibrils was then determined using fast-Fourier transform of the height profile of each fibril. For 2D FFT analysis, the fibril images were rotated with the straightened fibril axis aligned vertically. The images were subsequently zero-padded to squares prior to 2D FFT. All data analyses were performed using Matlab (MathWorks, Natick, Massachusetts)

## AUTHOR CONTRIBUTIONS

L.L. designed the research, developed the analytical software tools, analysed the data and drafted the manuscript. C.J.S. and M.F.T. designed the research and analysed the data. L.C.S. designed the research, provided assembly reagents and methods, and analysed the data. W.F.X. designed the research, developed the analytical software tools, analysed the data, managed the research, and drafted the manuscript. The manuscript was edited through contributions of all authors.

## ACKNOWLEDGEMENTS

We thank the members of the Xue, the CJ Serpell, and the LC Serpell groups, as well as the Kent Fungal Group for helpful comments throughout the preparation of this manuscript. We thank Ben Blakeman for acquiring and sharing the image data used in this report. This work was supported by funding from Biotechnology and Biological Sciences Research Council (BBSRC), UK grant BB/S003312/1, as well as Engineering and Physical Sciences Research Council (EPSRC), UK DTP grant (EP/R513246/1 for L.L).

